# Epistasis between SARS-CoV-2 M and N Proteins Balances Particle Assembly and Immune Evasion

**DOI:** 10.64898/2026.06.29.735107

**Authors:** Aldo Barrera-Vasquez, Mir M. Khalid, Hade Ramos, Julia Rosecrans, Marcela Ferres, Jenniffer Angulo, Melanie Ott, Taha Y. Taha

## Abstract

Since its emergence in the human population, SARS-CoV-2 has continuously evolved to evade immune responses and robustly establish global circulation. In this process, the structural viral membrane (M) protein has accumulated amino acid changes whose impact on viral particle assembly and innate immune evasion remains incompletely understood. Here, we designed a SARS-CoV-2 replicon system lacking M that assesses the influence of transiently transfected M protein variants on viral particle production independently of viral RNA replication. We found that M protein variants have reduced particle assembly while innate immune antagonism functions are strengthened. Notably, the assembly defect is rescued by co-evolving N protein variants, highlighting how SARS-CoV-2 evolution coordinates between two of its structural proteins to optimize viral infection. Our work underscores the complex evolutionary trajectories of SARS-CoV-2 variants across different viral proteins and informs future therapeutic strategies targeting viral assembly and limiting infection.

**Author summary:** Since the onset of the COVID-19, SARS-CoV-2 has been changing continuously in ways that help it spread efficiently and evade human defenses. The viral membrane (M) protein is a key structural component required to form new viral particles. Nevertheless, it has accumulated changes over time with overlooked functions. In this study, we developed an experimental system to examine the impact of these changes on virus assembly and host immune system. We found that recent M variants are better than older variants at suppressing immune responses but less efficient at forming viral particles. This defect seems to be compensated for by coordinated changes in another structural protein, nucleocapsid (N). These findings reveal that SARS-CoV-2 evolution involves trade-offs between different viral characteristics, with changes in one protein balancing changes in another. Understanding these trade-offs provides new insight into how the virus adapts to humans and may help guide therapeutic strategies to limit infection.

## Introduction

Since the onset of the COVID-19 pandemic, overall transmission levels of SARS-CoV-2 and the incidence of severe disease have declined [1, 2]. Nevertheless, SARS-CoV-2 continues to circulate worldwide. National and regional genomic monitoring systems reveal a complex landscape of Omicron-derived lineages with recombinant variants circulating at different frequencies and reflecting a dynamic process of adaptation to their hosts [3]. Recombinants such as XFG have accounted for a substantial proportion of recently sequenced cases in most countries, alongside other sub-lineages like NB.1.8.1 and BA.3.2 [2, 4]. Although updated vaccines continue to provide sufficient protection against severe disease, the ongoing evolution of SARS-CoV-2 underscores the need for sustained testing, genomic surveillance, and immunization efforts to mitigate its public health impact.

SARS-CoV-2 variants have been primarily classified based on mutations in the spike (S) protein, which mediates viral entry and is the main target of neutralizing antibodies [5]. Recent evidence suggests that SARS-CoV-2 evolution has shifted from an initially divergent pattern that gave rise to different variants of interest (VOI), towards a more constrained, convergent pattern of adaptation, with a reduced effective rate of variability in S [6–8]. For instance, extant Omicron strains share conserved mutations within the S receptor-binding domain (RBD), such as Q498R and N501Y, which preserve high-affinity binding to the SARS-CoV-2 cellular receptor ACE2 while compensating for mutations that promote antibody escape, including substitutions at positions 484-486 [9]. Beyond the S protein, other structural viral proteins, including the nucleocapsid (N), envelope (E), and membrane (M) proteins, exhibit lower overall variability but show fixation of a limited number of substitutions that may contribute to viral fitness, host immune modulation, and disease outcome [10]. Given the extensive focus on S protein evolution, the functional consequences of mutations in other structural proteins remain comparatively understudied.

The SARS-CoV-2 M protein, the most abundant structural component in the virion, comprises 222 amino acids organized into four domains: a short disordered N-terminal ectodomain (residues 1-18), three transmembrane helices (TM1, TM2, and TM3; residues 19-105), a hinge region (residues 106-116), and a C-terminal intravirion β-sheet domain (residues 117-201) [10, 11]. M is a multifunctional protein that plays a central role in virus assembly, morphology, and replication, orchestrating virion formation through interactions with S, E, and the N protein bound to genomic RNA (viral ribonucleoprotein complex, vRNP). These interactions coordinate genome packaging into budding particles and contribute to shaping the viral envelope. In addition to its structural functions, M modulates host cell responses by suppressing immune signaling pathways such as interferon production, thereby enhancing viral replication [11]. Throughout SARS-CoV-2 evolution, the M protein has accumulated a limited number of recurrent mutations, several of which reached frequencies above 1% within Omicron lineages. These include several exclusive substitutions at position three (D3G in BA.1, D3N in BA.5, and D3H in JN.1), as well as Q19E and A63T (BA.1 and beyond), and T30A and A104V (JN.1). Notably, D3H and the mutations located in the transmembrane domain of M remain conserved among the currently predominant circulating Omicron sub-lineages [11, 12]. The persistence of these mutations suggests a pleiotropic evolutionary benefit, but their specific impact on assembly and immune modulation remains poorly understood.

In this study, we developed a SARS-CoV-2 replicon system lacking M protein to investigate the role of M mutations in viral particle assembly independently of other functions that impact viral replication, such as immune evasion. We found that the M protein mutations in variants BA.1, BA.5, and JN.1 significantly impair viral assembly but exhibit enhanced innate immune evasion. These defects are compensated for by co-evolved N variants, a phenomenon of protein epistasis that restores efficient assembly and improves immune evasion. Our findings provide new insights into the coordinated evolution of SARS-CoV-2 structural proteins.

## Results

### An M-deficient SARS-CoV-2 replicon requires both exogenous M and N proteins to achieve robust viral particle assembly

To study the impact of M mutations on viral particle assembly and later infection, we developed a SARS-CoV-2 WA1 (the original 2020 isolate, herein referred to as “wild-type” or WT) M-dependent replicon by replacing the M open reading frame (ORF) with a sequence coding for the monomeric NeonGreen (mNG) fluorescent protein or secreted nanoLuciferase (Sec-nLuc) (Fig. 1A). The replicon plasmid was co-transfected with an M protein expression vector (pM) into BHK-21 producer cells to generate infectious particles (Fig. 1B). These particles are then used to infect receiver cells and reporter expression is measured as a surrogate for the efficacy of viral particle assembly (Fig. 1B). Because the replicon does not encode its own M RNA/protein, infection in the receiver cells cannot produce viral particles and therefore the infection is limited to a single round.

**Figure 1.**
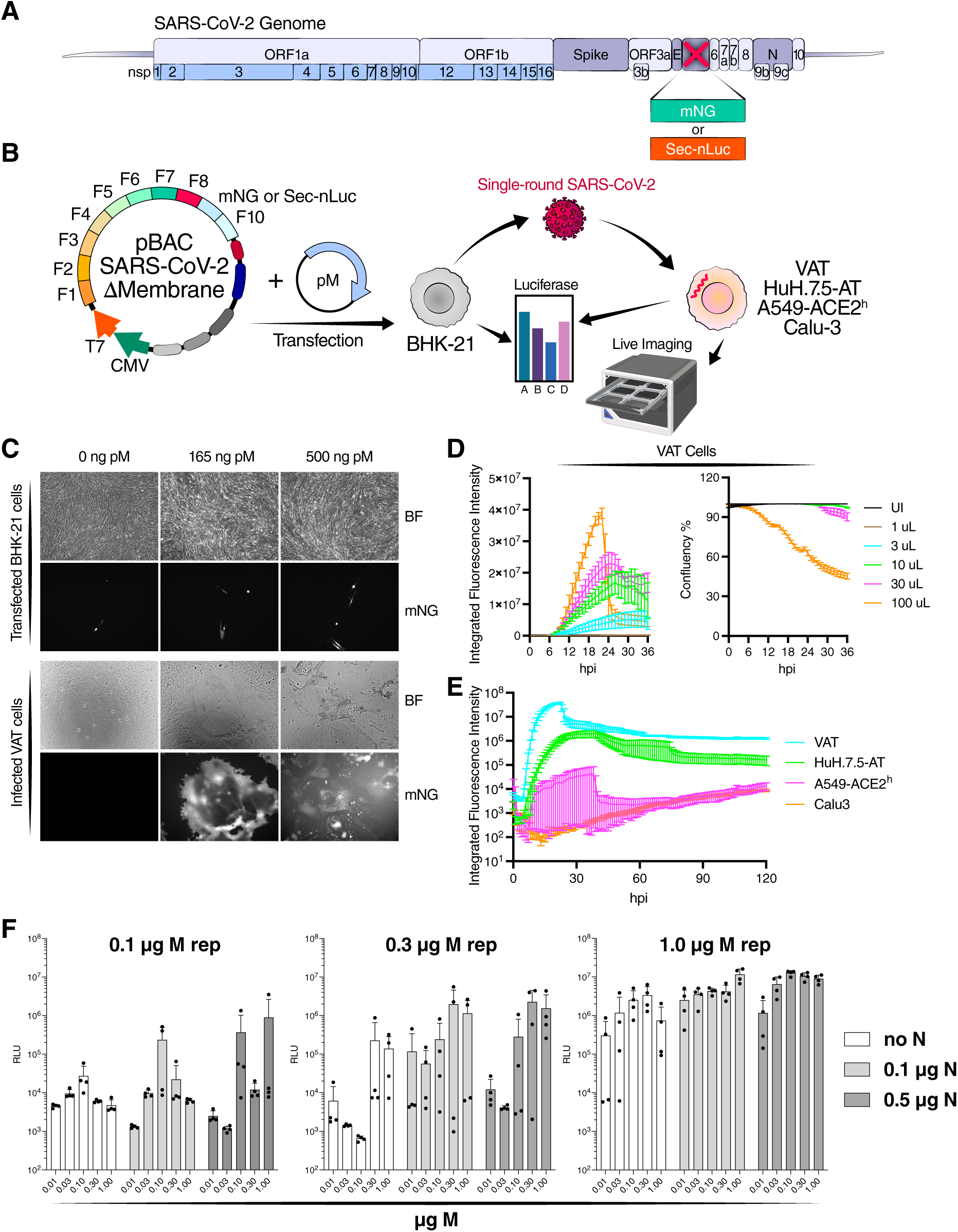
SARS-CoV-2 M replicon enables study of M protein role in assembly independent of viral replication. **A)** Schematic of the M replicon in the context of the SARS-CoV-2 genome. The M coding sequence (WA1 nt: 26,523-27,032) was replaced with either mNeonGreen (mNG) or secreted nanoluciferase (Sec-nLuc). **B)** Schematic representation of experiments employing the M replicon in either fluorescence or luciferase readouts. Viral particles produced by BHK-21 cells promote a single round of infection because they do not encode their own M protein and therefore cannot generate new infectious particles. **C)** Fluorescence readout of transfected BHK-21 cells and infected VAT cells 48 hours post-transfection or -infection. The amount of M expression vector (pM) used at the transfection step is indicated on the top portion of the panel. Images are representative of three independent biological replicates. Syncytium is observed as a rounded pattern of fluorescence. **D)** Live imaging readout and confluency of infected VAT cells with indicated volume of supernatant containing single-round particles. **E)** Live imaging readout of infection of different cell lines from the experiment in (C). All cells were infected with 100 μL of supernatant containing single-round particles. Data in (D) and (E) are presented as mean +/- SD of two technical replicates. **F)** Luciferase readout of infected VAT cells with indicated amounts of M replicon and M and N expression vectors. Data are presented as mean +/- SD of four independent biological replicates.

To demonstrate the dependence of this system on M protein for viral particle assembly, we co-transfected the SARS-CoV-2 mNG M-replicon into cells along with varying amounts of pM before harvest and infecting receiver cells (Fig. 1C). While fluorescence in transfected cells was relatively constant across all samples and was not dependent on M expression, fluorescence in infected VAT cells increased with higher amounts of transfected pM, suggesting increased particle production without effects on viral RNA replication and translation. Increased fluorescence was accompanied by increased syncytia formation (Supplementary Video S1) reflecting S protein expression on the surface of infected cells promoting cell-to-cell fusion via overexpressed ACE2 and TMPRSS2.

To optimize our assay, we modified a number of variables. First, we wanted to test of reporter activity in receiver cells correlates with the amount of inoculum. We tested various amounts of producer cell supernatant in VAT cells, and measured both fluorescence intensity and cytotoxicity, as measured by cell confluency. As expected, higher volumes of the supernatant containing the inoculum particles were associated with higher reporter activity and cytotoxicity in VAT cells (Fig. 1D). Therefore, reporter expression in the receiver cells can be used as a surrogate for the efficacy of the initial particle assembly. Next, we infected different cell lines and monitored fluorescence kinetics over 120 hours post-infection. We chose standard cell lines in SARS-CoV-2 research, including VAT (monkey kidney), HuH.7.5-AT (human liver), A549-ACE2^h^, and Calu3 (human lung). Fluorescence peak intensity was highest in VAT cells (∼10^7^ RLU) followed by HuH.7.5-AT cells (∼10^6^ RLU), A549-ACE2h ((∼10^4^ RLU), and Calu3 cells (∼10^4^ RLU).

All cell lines reached peak fluorescence intensity around 24 hours post-infection, except for Calu3 cells, which continued to increase over the course of the experiment. Therefore, we continued using VAT cells for subsequent experiments given the high infection efficiency and reporter expression in these cells. Finally, we modified the amount of the transfected plasmids M replicon (0.1-1 µg) and pM (0.01-1 µg) in the presence or absence of pN (0-0.5 µg), a plasmid encoding the SARS-CoV-2 WT Nucleocapsid protein. We reasoned that providing N in trans should help launch viral replication and assembly as previously reported [13]. While all experimental conditions resulted in a detectable luciferase signal in receiver cells, a high amount of the replicon (1 µg) plasmid was necessary to achieve high signal (>10^6^ RLU) and low variability (Fig. 1F). As expected, increasing amounts of pM enhanced the luciferase signal but the signal reached a plateau after 0.3 µg in most cases (Fig. 1F). The presence of pN was critical for efficient and consistent luciferase signals (Fig. 1F). Therefore, both medium-to-high levels of M and N proteins are necessary for viral particle assembly in this replicon system. Interestingly, the luciferase signal in transfected and infected cells negatively correlated with the amounts of pM (Supplementary Fig. 1), suggesting that higher assembly efficiency at higher pM amounts might be depleting viral transcripts in transfected cells, thereby reducing reporter expression. Collectively, these data demonstrate the utility of this replicon system in assessing the role of M in viral particle assembly.

### M proteins from the BA.1, BA.5, and JN.1 variants support less particle assembly than wild-type M protein

Next, we used our optimized assay conditions to test the impact of Omicron M variants on viral particle assembly in combination with various N Omicron variants (Fig. 2A and 2B). First, we assessed the effect of growing amounts of each M variant in combination with a fixed amount of wild-type N to account for potential differences in M expression levels. Only the BA.2 M variant yielded a dose-response curve similar to that of WT M. By contrast, the BA.1, BA.5, and JN.1 M variants produced a significantly reduced luciferase signal compared with wild-type M across all pM amounts (Fig. 2C). To determine the contribution of each M mutation to this reduced assembly phenotype, we constructed pM constructs each carrying a single amino acid change found in these variants (Fig. 2B) and tested their impact on viral particle assembly. Only D3N, D3H, and T30A yielded a reduced luciferase signal compared to wild-type M and only at lower pM amounts (Fig. 2C). By contrast, Q19E increased the luciferase signal significantly compared with wild-type M protein (Fig. 2C).

**Figure 2.**
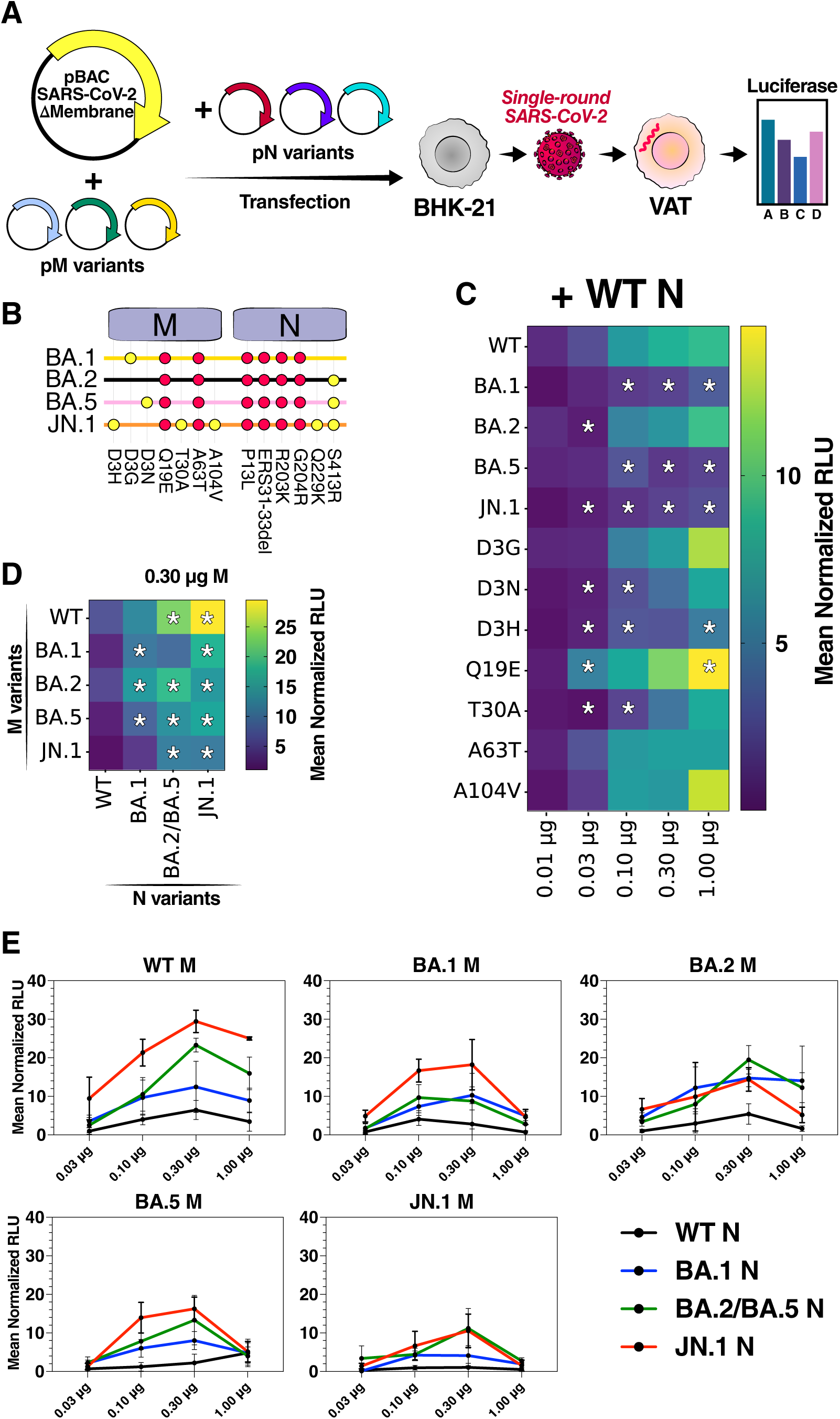
Reduced ability of M protein variants to produce infectious particles is rescued by co-evolved N variants in the replicon assay. **A)** Schematic representation of experiments to determine the impact of M and N protein amino acid changes on viral particle assembly. **B)** Overview of M and N protein amino acid changes throughout SARS-CoV-2 Omicron evolution. **C)** Luciferase readout of infected VAT cells normalized to 0.10 μg WT M protein of indicated M protein variants at various plasmid amounts. **D)** Luciferase readout of infected VAT cells normalized to WT M and N proteins. **E)** Luciferase readout of infected VAT cells normalized to 0.03 μg M protein. For all data in (D-F), M replicon and N expression vectors were fixed at 1 μg and 0.5 μg, respectively, and the data are presented as mean +/- SD of four independent biological replicates. *, p < 0.05 using two-sided Student’s T-test.

Strikingly, the combined effect of single amino acid changes does not explain the observed effect of M variants. These findings suggest strong intragenic epistatic interactions within the M protein. For instance, all amino acid substitutions found in BA.1 M (i.e. D3G, Q19E, and A63T) have individual effects on assembly that mirror or exceed those of wild-type M, while BA.1 M allows significantly less assembly than wild-type M (Fig. 2C). The discrepancy is less striking for BA.5 M and JN.1 M because at least one of their single amino acid changes yields a reduced luciferase signal (i.e. D3N for BA.5 and D3H/T30A for JN.1). The lack of correlation between the phenotypes of M variants and single amino acid substitutions suggests epistatic effects between the individual mutations. Overall, these data demonstrate that in the context of the WT SARS-CoV-2 genome, M variants BA.1, BA.5, and JN.1 have a significantly reduced ability to assemble viral particles and that substitutions in BA.1 M exert negative epistasis on each other.

### N protein variants rescue the particle-assembly deficiencies of M protein variants

The reduced viral particle assembly of BA.1, BA.5, and JN.1, despite their high transmission and replication in the population, prompted us to investigate potential compensating mechanisms. We focused on the N protein since it is known to interact with M [14–16], and we had already shown that adding N protein to our assay boosts particle formation (Fig. 1F). We generated constructs expressing the N proteins of the BA.1, BA.2/BA.5, and JN.1 variants, and combined them to various amounts of the variant pM constructs (Fig. 2A and 2B). We found that all N variants increased viral particle assembly in the presence of almost all M variants (Fig. 2D and 2E). The JN.1 N variant had the highest impact and rescued all M variants to almost equal levels (Fig. 2E). For instance, when 0.30 µg of pM were used, all N variants increased luciferase signal compared with wild-type N for all tested M variants, with some exceptions that did not reach statistical significance (Fig. 2D). This general enhancement of particle assembly suggests that N variants evolve toward an optimization of their viral particle assembly phenotype, consistent with our previous observations [17].

### M protein variants have reduced VLP production that is rescued by co-evolved N variants

To determine whether compensatory effects on particle assembly are limited to M and N proteins, we employed virus-like particles (VLPs) that are composed of all four structural proteins and the authentic SARS-CoV-2 packaging signal [18]. We constructed VLP plasmids of B.1 (wild-type), Delta, and all Omicron structural protein variants until JN.1 (Fig. 2B) [19], and transfected them into HEK293T cells to produce VLPs. We purified the VLPs by ultracentrifugation and determined their abundance by Western blot analysis for the Spike and N proteins (Fig. 3A). Strikingly, all Omicron variant VLPs showed similar abundance to wild-type VLPs (Fig. 3B). By contrast, the Delta variant VLPs were more abundant, as we have previously reported [20]. The behavior of the Omicron VLPs may reflect the compensatory effect of N mutations on M mutations we observed in our replicon assay. Alternatively, there might be other compensatory effects involving E and S. These findings further support the importance of genetic context in determining structural protein phenotypes.

**Figure 3.**
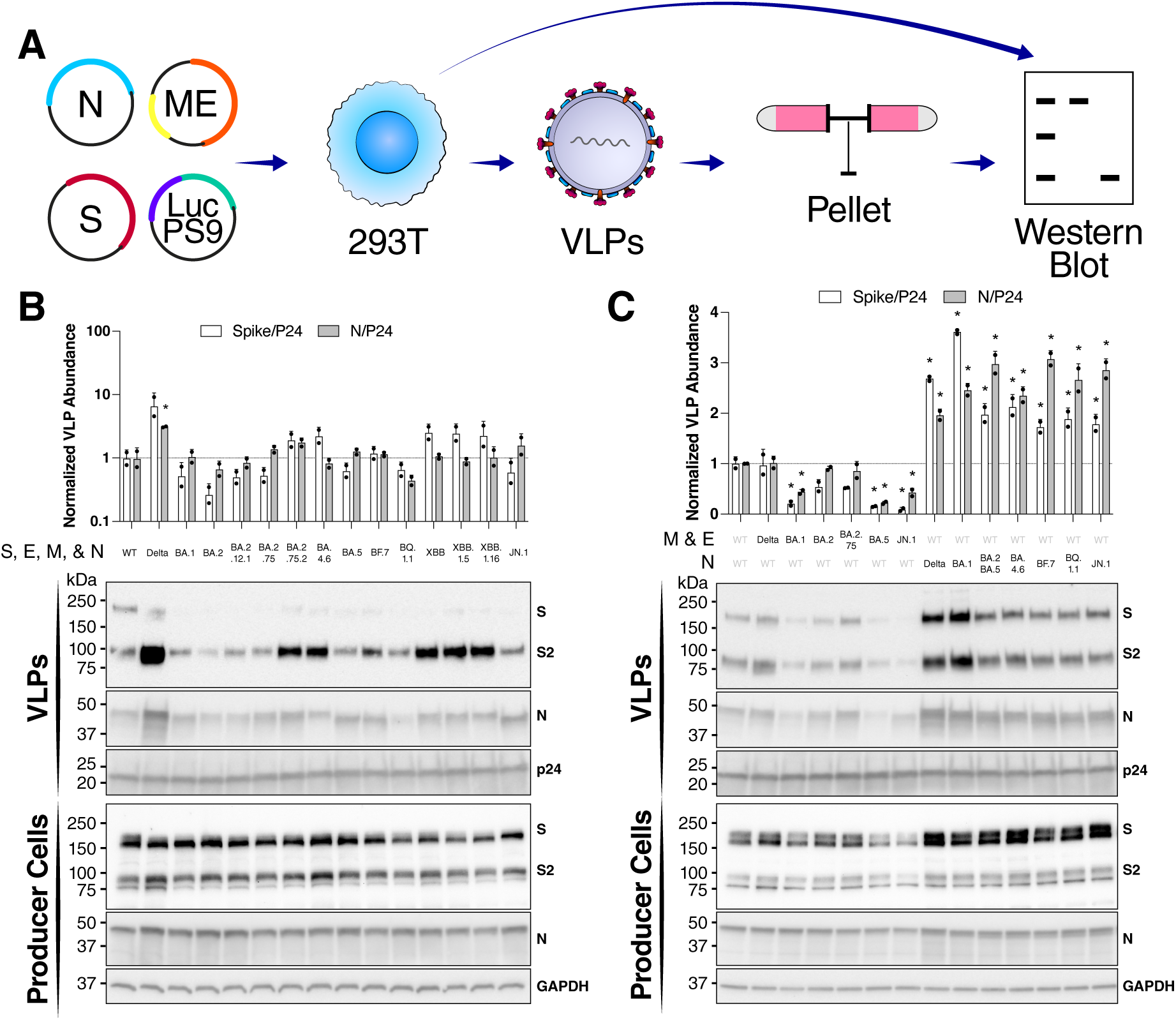
Reduced ability of M protein variants to produce infectious particles is rescued by co-evolved N variants in the VLP assay. **A)** Schematic representation of VLP production, purification, and analysis experiments. **B and C)** Western blot analysis of VLP producer cells and purified VLPs for the Spike and N proteins. GAPDH was used as a cellular internal control, and p24 was used as VLP purification internal control for spike-in lentiviral particles. VLPs comprising all co-evolving structural proteins are shown in (B), and VLPs with variable M/E or N proteins are shown in (C). Blots are representative of two independent biological replicates. The top panel shows VLP abundance based on the normalized Spike or N proteins as mean +/- SD. *, p < 0.05 using two-sided Student’s T-test.

To distinguish these hypotheses, we made VLPs with individual structural protein variation in the context of wild-type VLPs (Fig. 3C and supplementary Fig. 2). Altering the S protein in the context of wild-type M, E, and N proteins did not impact VLP assembly (supplementary Fig. 2). However, VLPs bearing the BA.1, BA.5, and JN.1 M proteins in the context of wild-type N and S were significantly less abundant than those bearing wild-type M protein (Fig. 3C). Of note, all Omicron variants studied here have a single amino acid substitution in E protein T9I, except for BA.2.75- and XBB-derived lineages that have an additional T11A mutation [19]. However, the E mutations were not associated with any impact on VLP assembly (compare WT, BA.1, BA.2, and BA.2.75 VLPs in Fig. 3C). N variants in the context of wild-type M, E, and S all had enhanced VLP assembly compared with wild-type N (Fig. 3C). These data show that consistent with the replicon studies, the assembly phenotype of M variants depends on the context of other structural proteins in the SARS-CoV-2 virion.

### Evolved M protein variants escape innate and adaptive immune mechanisms

Viral protein evolution typically balances different functions [17], and we wondered whether the reduced viral particle assembly is the cost M proteins pay to optimize other functions. Previous reports have shown that the SARS-CoV-2 M protein antagonizes the innate immune system [21–24], including the mitochondrial antiviral signaling (MAVS) pathway. To test M proteins’ effects on these systems, we combined an interferon beta luciferase (IFN-β-Fluc) reporter with three key activators of this pathway: MAVS, the retinoic acid-inducible gene I (RIG-I), and the TANK-binding kinase 1 (TBK1). We expressed the Influenza A Virus (IAV) NS1 protein, a known inhibitor of the IFN-β response, as a positive control [25] and the Sin Nombre Virus (SNV) N protein, which is not known to affect IFN-β, as a negative control [26]. We observed that wild-type M inhibited IFN-β-Fluc activation by MAVS, RIG-I, and TBK1 to a minimum of around 50% in an M-dose-dependent fashion (Fig. 4B). Then, we transfected each activator with M variants and measured IFN-β activation. M protein levels did not differ significantly across all tested variants after normalization to the endogenous glyceraldehyde-3-phosphate dehydrogenase (GAPDH) levels (Fig. 4D). The M protein of Omicron variants BA.1 and JN.1 antagonized MAVS’s activation of IFN-β-Fluc better than wild-type M protein did (Fig. 4C). BA.5 M had similar inhibitor activity and BA.2 M had reduced inhibitor activity compared with wild-type M (Fig. 4C). Of the single amino acid substitution variants, only T30A M enhanced the inhibitory effect of M protein, while all other substitutions had reduced inhibitor activity (Fig. 4C). BA.1 M also antagonized RIG-I and TBK1 significantly more than did wild-type M, but the other natural variant M proteins did not, and none of the single substitution variants did either (Fig. 4C). In fact, BA.2 M and most single-substitution M proteins showed a reduced ability to antagonize TBK1 (Fig. 4C). The discrepancy between the inhibitory activity of BA.1 M and that of M proteins with the single amino acid substitutions carried by BA.1 M suggests an epistatic effect consistent with the effect observed with viral assembly in our replicon assay (Fig. 2C). Collectively, these data suggest an enhancement in innate immune antagonism of BA.1, BA.5, and JN.1 M protein variants. These findings provide additional evidence of intragenic epistasis within the M protein, extending beyond assembly phenotypes to innate immune antagonism.

**Figure 4.**
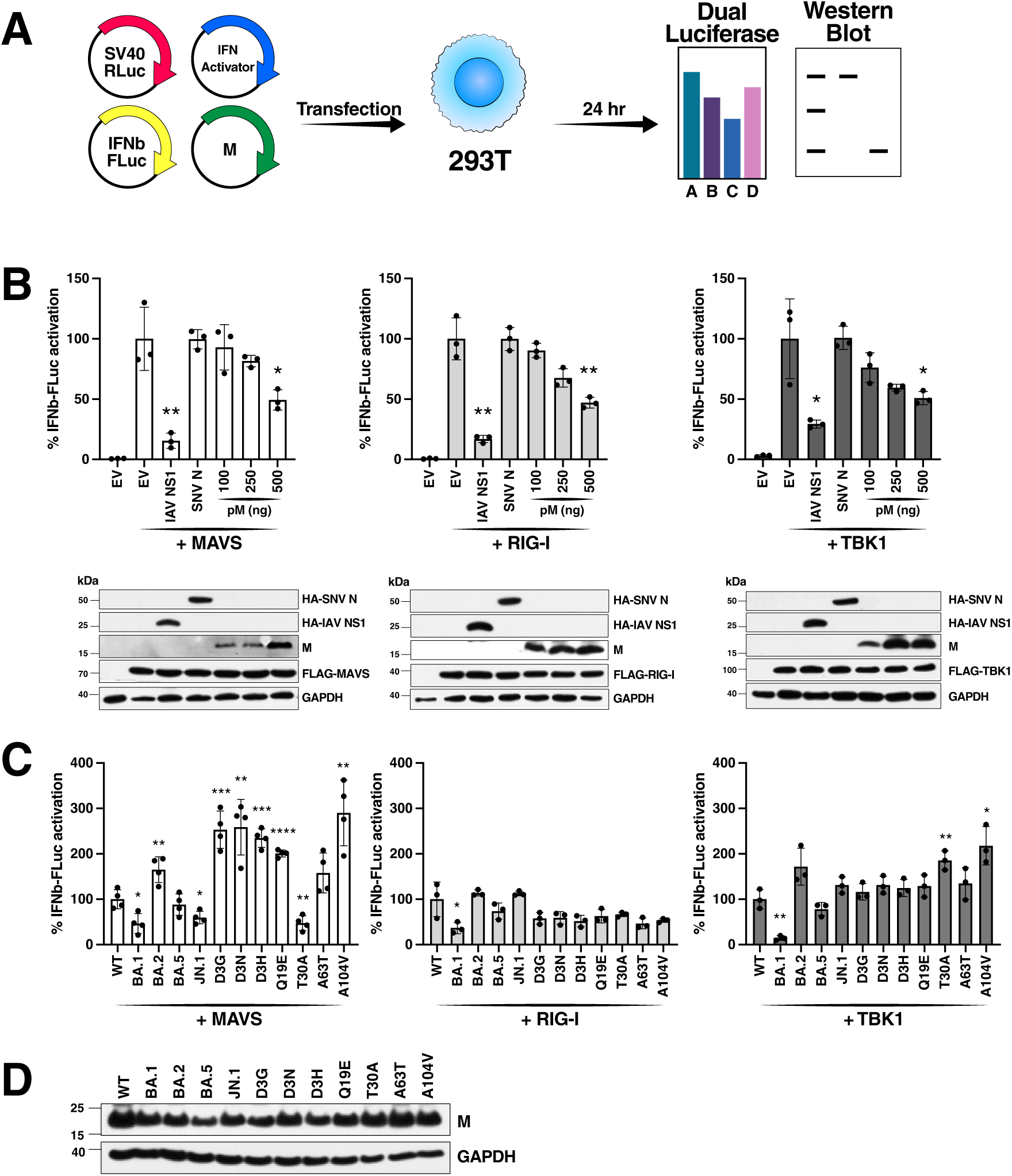
M protein variants antagonize innate immunity. **A)** Schematic representation of the innate immune inhibition assays. The Renilla luciferase (RLuc) construct is cotransfected as an internal control. Firefly luciferase (FLuc) measurements from the interferon beta luciferase (IFN-β-Fluc) reporter are divided by the internal control RLuc measurement (i.e. dual luciferase readout) to account for any transfection variability or cell loss during transfection. **B)** Luciferase readout and Western blot analysis of cells transfected with indicated innate immune activators and ng amount of M protein expression vector. Influenza virus (IAV) NS1 and Sin Nombre virus (SNV) N were utilized as positive and negative controls, respectively. Data are presented as mean +/- SEM of three independent biological replicates and normalized to vector control. Blots are representative of three independent biological replicates. **C)** Luciferase readout and Western blot analysis of cells transfected with indicated innate immune activators and M protein variants. Data are presented as mean +/- SD of at least three independent biological replicates normalized to the WT M protein. Blot is representative of three independent biological replicates. *, p < 0.05; **, p < 0.01; ***, p < 0.001; ****, p < 0.0001 by two-sided Student’s T-test.

M protein has also been reported to elicit antibody responses to its N-terminal portion [27–29], which is located on the outer side of the virion membrane (aa 1-18). Therefore, we asked whether substitutions in position 3 of the M protein that are occurring in natural variants (Fig. 2B) could evade antibody responses. We developed an ELISA assay with biotinylated versions of the 20-amino acid N-terminal ectodomain carrying the wild-type sequence, BA.1 substitution D3G, BA.5 substitution D3N, and JN.1 substitution D3H (Fig. 5A). The peptides were immobilized onto streptavidin-coated plates and subsequently incubated with 32 sera samples from a cohort of first-wave COVID-19 infected individuals [30]. A horseradish peroxidase (HRP)-conjugated secondary antibody allowed us to quantify the binding of the serum antibodies to the peptides by measuring the OD_450_. We used a peptide corresponding to amino acids 153-170 of N as a control for non-S antibody responses [28]. Anti-N antibody titers measured were higher than anti-M antibody titers, and a greater number of sera were anti-N positive than anti-M positive (24/32 vs. 10/32; Fig. 5B). Moreover, the sera’s antibody titers were lower against M peptides containing mutations D3G, D3N, and D3H than against WT M. These data suggest that M protein evolution could potentially introduce substitutions at position 3 to evade antibody recognition in previously infected individuals.

**Figure 5.**
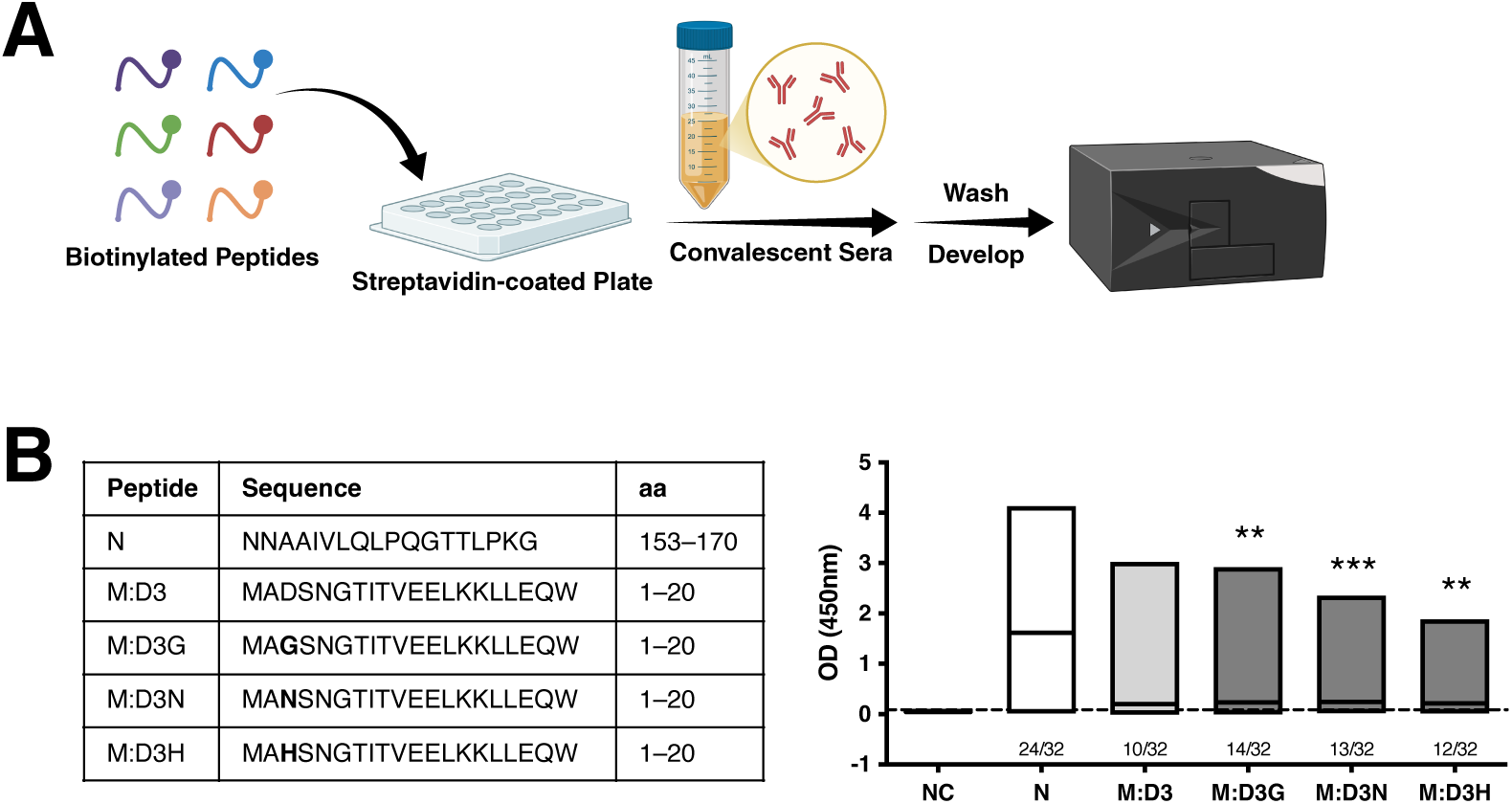
M protein variants at position 3 evade antibody recognition. **A)** Schematic representation of ELISA to detect human antibody binding to M protein peptides. **B)** The table on the left shows the sequence of the tested peptides and their amino acid position in the M and N proteins. The graph on the right shows the optical density at 450 nm (OD_450nm_) of ELISA for human IgG using indicated peptides as the antigen. The data are plotted as floating bars (min to max) and the mean OD_450nm_ measured for 32 serum samples. The number of samples above the detection limit is indicated under each box. NC indicates negative control peptide. **, p < 0.01; ***, p < 0.001 by a two-sided Wilcoxon’s T-test.

## Discussion

Since the emergence of SARS-CoV-2 in humans, the virus’s M protein has accumulated far fewer amino acid substitutions than the S protein, suggesting that M is constrained by a stringent purifying selection that preserves its essential roles in virion assembly and structural integrity. Here we present evidence that in fact, M protein evolution within the Omicron lineages has compromised viral particle assembly, but that they also conferred protection against the host’s immune response. Moreover, we demonstrate that M and N proteins have co-evolved in Omicron lineages such that N variants compensate for M variants’ defects in particle assembly. Finally, we uncover evidence of epistatic relationships between sequence variants within the M proteins. Our findings illustrate the usefulness of our combined new M-dependent replicon and VLP systems to monitor the impact of structural protein variants in the context of more than one viral protein at a time. Such monitoring is crucial for the proper surveillance of emergent SARS-CoV-2 and other coronaviral variants.

Our data suggest that M protein evolution trades losses in viral particle assembly for gains in immune evasion. We have previously shown that Omicron BA.1 M protein has lower particle assembly compared with the B.1 and Delta variants [20]. Here we show post-BA.1 M proteins, except for BA.2, have significantly reduced viral particle production in both the replicon and VLP assays. Previous studies have suggested that mutations in the TMH3 might impact M dimerization and M-E interaction [31, 32], however, mutations A63T and A104V do not show an apparent effect on viral particle assembly. In terms of innate immune antagonism, M protein has been shown to inhibit the MAVS pathway as a common mechanism for coronaviruses to compromise type-I IFN responses. Both SARS-CoV and Middle East respiratory syndrome coronavirus (MERS-CoV) M proteins inhibit RIG-I and the TBK1-dependent activation of IFN regulatory factor 3 (IRF3) [33, 34]. The SARS-CoV-2 M protein promotes the ubiquitination and degradation of TBK1 [21, 22]. Here we reproduce the inhibitory effect of M protein on MAVS, RIG-I, and TBK1, and find that the BA.1 and JN.1 M variants, and to a lesser extent BA.5 M, improve the inhibitory activity of M protein on MAVS-induced IFN-β response compared with wild-type M. BA.1 M protein also inhibits both RIG-I- and TBK1-induced IFN-β response compared with wild-type M, showing a broader antagonism of upstream type I interferon signaling pathways. This pattern aligns with the hypothesis that Omicron BA.1 may have emerged under strong within-host immune pressure in an immunocompromised individual [35], where selection could favor variants capable of suppressing multiple innate sensing nodes, including RIG-I/MDA5–MAVS signaling, while subsequent Omicron derivatives appear to refine this strategy toward more specific targeting of MAVS-dependent antiviral signaling. These data are also consistent with systems virology approaches that have shown that BA.2 infection induces higher level of innate immune responses compared with ancestral SARS-CoV-2, BA.1, or BA.5 infections [7]. Fu et al. showed that the ectodomain TMH1 and TMH2 regions are essential for MAVS inhibition [22], but our data show that only T30A, but not Q19E and A63T, enhance M’s inhibitory function. Previous studies have also suggested that mutations in the TMH3 might impact M dimerization and M-E interaction [31, 32], however, mutations A63T and A104V do not show an apparent effect on viral particle assembly. Collectively, our study highlights how M protein evolution within Omicron lineages optimized its innate immune antagonism functions at the cost of viral particle assembly.

Viruses often co-evolve compensatory mechanisms to enable risk-taking mutations elsewhere in the viral genome. Here we demonstrate an epistatic relationship of M and N proteins in Omicron lineages such that N variants compensate for M variants’ defects in particle assembly. One of the critical determinants of coronavirus assembly is the interactions of the M protein with vRNP complexes, containing N proteins and viral RNA [11, 16]. We previously showed that substitutions in the N protein linker region of the Omicron BA.1 variant enhance viral RNA packaging and particle production [18, 20]. Here we show that post-BA.1 N substitutions in the linker region (Q229K) as well as C-terminal domain (S413R) enhance viral particle production. This enhancement would maintain an overall robust viral particle assembly for infection and transmission, despite disadvantageous substitutions in the M protein. Indeed, we find that recent N protein variants, such as JN.1, rescue the assembly defect of all natural M protein variants. Our data therefore suggests that N evolution is endowing a broad assembly-promoting effect rather than a specific effect restricted to cognate M-N pairs.

Our results indicate significant epistasis within the M protein that likely governs its evolution. We find single amino acid substitutions have drastically different assembly phenotypes in different M protein contexts. For example, substitutions D3G and D3N decrease particle assembly of the BA.1 and BA.5 variants (i.e. in combination with Q19E and T30A), respectively, although they have no impact on particle assembly in a wild-type M background. Similar to the assembly phenotype, there was significant epistasis with regards to M protein’s role in innate immune antagonism. We observed that all individual substitutions, but T30A, revert the inhibitory effect on MAVS but not on RIG-I or TBK1. Since the BA.1, BA.5, and JN.1 M variants promote the inhibitory role of M, but BA.2 does not, indicate that substitutions in position 3 enhance MAVS antagonism only in the presence of Q19E and A63T, but not WT background. These findings are consistent with reports of significant epistasis in SARS-CoV-2 proteins [17, 36–38], especially the Spike protein [39, 40]. However, this is the first time that compensatory effects of M protein substitutions are described. These observations suggest that the functional consequences of M mutations cannot be inferred from individual substitutions alone, highlighting the importance of genetic context in shaping viral phenotypes.

Substitutions in M do not appear to induce structural changes that may impact other M protein properties. To determine whether M protein WT, BA.1, BA.2, BA.5, and JN.1 variant structures differ, we predicted homodimer structures of each variant using AlphaFold 3. We found that they are all structurally indistinguishable at the backbone (Cα) level (≤1.1 Å) after accounting for the AF3 sampling noise (noise floor ∼1.65 Å) within each variant’s 5-model ensemble (Supplementary Fig. S3). We did not find any residues, including the five mutated positions, that show a variant-specific conformational change. It is important to note a limitation of our analysis as we only considered conformational change at the Cα-backbone level and not fully encompassing sidechain/packing effects of the substitutions. Collectively, the effects we observe on virion assembly and innate immune antagonism may be due to subtle rather than large-scale structural alterations.

M protein substitutions impact its recognition by the adaptive immune system. The immunogenicity of M has been reported to be comparable with S and N, finding high ectodomain-specific (aa 1-18) IgG titers in convalescent and vaccinated individuals [28]. Previous studies have shown that antibodies against non-S structural proteins do not induce SARS-CoV-2 neutralization but promote the antibody-dependent cell cytotoxicity (ADCC) induced by Fc receptor-expressing cells, limiting viral replication [29, 41–43]. Since the emergence of Omicron, three substitutions have been observed within the M protein ectodomain: D3G in BA.1, D3N in BA.5, and D3H in JN.1. Recently, however, the BA.3.2 variant has again reverted to wild-type at position 3 [4]. This recurrent evolution at position 3 suggests the presence of an ongoing selective pressure at the N-terminal region of M, and our results show that limited sequence variation may have important functional consequences. We observed that these substitutions reduce binding by antibodies present in sera from critically ill patients, suggesting a potential role for these recurrent mutations in immune evasion. It is important to note that a limitation of our study is that a limited number of sera exhibited detectable reactivity and only few showed high anti-M titers, indicating that this finding requires further validation using larger cohorts, ectodomain-specific monoclonal antibodies, and functional ADCC assays.

Our study extends the many instances of evolutionary tradeoffs between structural integrity and immune evasion for SARS-CoV-2. A well-known example involves substitutions in antigenic sites of the Spike protein that reduce its stability and affinity to the ACE2 receptor, resulting in lower cell entry efficiency, but conferring enhanced evasion from neutralizing antibodies. These substitutions thereby allow the virus to persist and transmit in populations with pre-existing immunity from vaccination or prior infection [5]. Another example of a functional evolutionary tradeoff is the accessory protein ORF8, which restricts cell and virion surface expression of S protein, thereby reducing viral particle infectivity but limiting infected cell recognition by antibodies and cytotoxic immune cells [44, 45]. Throughout the pandemic, SARS-CoV-2 evolution tuned the expression of ORF8 in variants of concern by introducing premature stop codons [46].

Molecular tools to study coronaviral evolution help pinpoint specific viral processes under selection and improve monitoring strategies. Our new M-deficient replicon provides a unique opportunity to study the role of both M and N proteins in viral particle assembly independent of RNA replication. This is critically important as studying mutations in the full-length viral context may mask ongoing evolution for specific viral protein functions. The M-deficient replicon is incapable of producing viral particles unless M protein is provided in trans, and therefore it provides an excellent model for cell-to-cell spread independent of viral particle assembly or entry processes. This property also endows the system with high signal to noise compared with previous replicon systems and makes it useful as an antiviral testing platform at lower biocontainment settings.

All in all, we show that epistatic interactions between SARS-CoV-2 structural proteins and support a model in which mutations across the viral genome interact functionally to shape viral phenotypes (Fig. 6). Our work highlights that SARS-CoV-2 evolution involves trade-offs between different viral functions, with substitutions in one protein being balanced by changes in another. Understanding these interactions provides new insight into how the virus keeps adapting to humans and may help guide strategies to disrupt viral assembly as a therapeutic strategy to limit infection.

**Figure 6.**
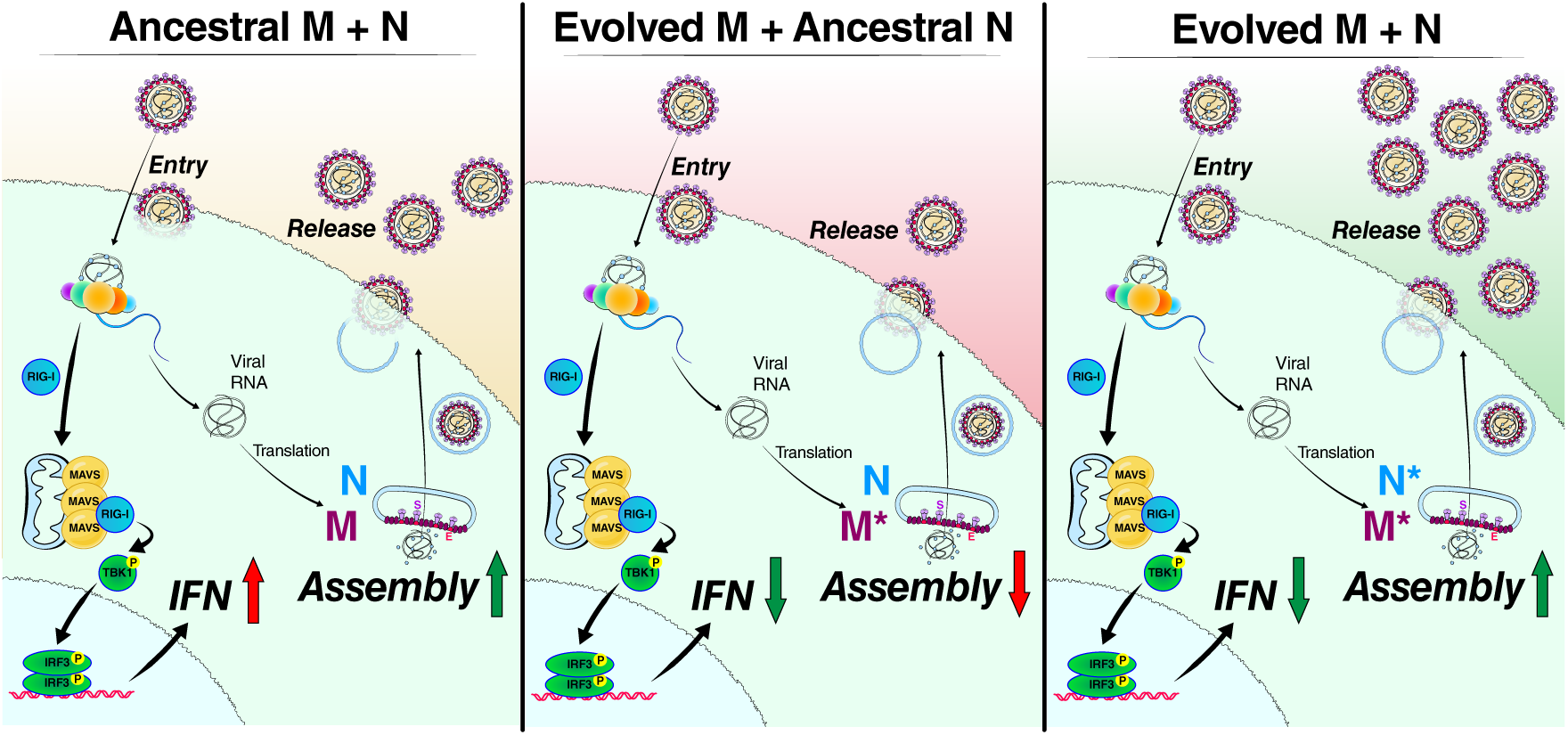
Proposed model for coordinated M and N protein co-evolution balancing innate immune antagonism and viral particle assembly. During infection, viral RNA replication is detected by the cytosolic sensors MDA5 and RIG-I, which signal through MAVS to promote TBK1 phosphorylation. Phosphorylated TBK1 translocate to the nucleus, where it activates IRF3 and drives type-I interferon (IFN) expression. In addition to its canonical role in virion assembly, the M protein suppresses innate immune signaling at multiple steps in this pathway. Over the course of viral evolution, M protein variants have increased innate immune antagonism, accompanied by a reduction in assembly efficiency. Compensatory evolution of the N protein restores viral particle assembly, functionally offsetting the assembly defects associated with evolved M variants.

## Materials and Methods

### Cells

BHK-21, 293T cells were obtained from ATCC and grown in standard media: DMEM (Corning) supplemented with 10% fetal bovine serum (FBS) (GeminiBio), 1x Glutamax (Corning), 1x non-essential amino acids (NEAA) (Corning), and 1x penicillin-streptomycin (Corning) at 37 °C, 5% CO_2_. A549 cells stably expressing high levels of ACE2 (A549-ACE2^h^) have been previously described and were maintained in standard media supplemented with 10 μg/mL blasticidin [45]. HuH.7.5 cells stably expressing ACE2 and TMPRSS2 (HuH.7.5-AT) have been previously described [17] and were maintained in standard media with 10 μg/mL blasticidin and 250 μg/mL hygromycin. Calu3 cells were obtained from ATCC and cultured in Advanced DMEM (Gibco) supplemented with 2.5% FBS, 1x GlutaMax, and 1x Penicillin-Streptomycin at 37°C and 5% CO_2_. Vero cells stably expressing human ACE2 and TMPRSS2 (VAT) (gifted from A. Creanga and B. Graham at NIH) were maintained in standard media with the addition of 10 μg/mL of puromycin. All cell lines are confirmed mycoplasma-free by quarterly testing.

### Plasmid Construction

Replicon plasmids were constructed as described previously [47]. Briefly, the M protein coding sequence (WA1 nt: 26,523-27,032) were replaced with the mNG2 or secreted nanoLuc coding sequences as described previously using NEB HiFi DNA assembly [48]. The 3’ region of the M coding sequence (WA1 nt: 27,033-27,191) was unaltered to preserve the TRS sequence of ORF6. All plasmids utilized for virus-like particle assays have been described previously [18] and mutations were introduced using NEB HiFi DNA Assembly [49]. The plasmids IFN-β–FLuc (p125-Luc), pCAGGs-FLAG-EV, pCAGGs-FLAG-RIG-I, pCAGGs-FLAG-MAVS, pCAGGs-FLAG-TBK1, HA-NS1 from IAV, and HA-SNV-N were previously described in [26] and kindly provided by Dr. Adolfo García-Sastre (Icahn School of Medicine at Mount Sinai, Nueva York, USA).

### Replicon Assays

Replicon assays were conducted as described previously with some modifications [19, 50, 51]. Briefly, 5×10^4^ BHK-21 cells were plated in 24-well plates. After 24 hours, the cells were transfected with different amounts of replicon (mNG or nLuc), M, and/or N plasmids (see figures) using X-tremeGENE 9 DNA Transfection Reagent at 1:3 ratio in serum-free Optimem and 1 hour incubation at room temperature (Millipore Sigma). The supernatant was replaced 4–6 hours post-transfection with 0.3 mL fresh medium. The supernatants were collected at 48 hours post-transfection and filtered using Pall 0.45 μm AcroPrep Advance Plate, Short Tip filter plates (polyethersulfone). The filtered supernatants were mixed with 2×10^4^ VAT, A549-ACE2^h^, HuH.7.5-AT, or Calu3 cells and plated in 96-well plates. For live imaging experiments, the cells were imaged every hour using an Incucyte S3 system. For luciferase experiments, six hours post-infection, the cells were washed once with 300 μL PBS and 100 μL of fresh medium was added. Luciferase measurements in the supernatants were conducted at 24 hours post-infection using 50 μL of supernatant and 50 μL of nano-Glo luciferase reagent (Promega).

### Virus-like particle Western blot

293T cells were seeded at 7×10^6^ cells per 15-cm dish and transfected the next day (50% confluent) with 6.64 µg pMIRESE, 13.32 µg pN R203M, 20 µg pLuc PS9, and 0.4 µg pSpike variant plasmids using Xtremegene 9 transfection reagent (Sigma Aldrich) (1:3 ratio). The media was changed 12-16 hours post-transfection. After 48 hours, the supernatant was collected and 0.45 µm filtered. The cells were washed with PBS and collected in RIPA lysis buffer and stored at −20 C. 1 mL of lentivirus preparation was added to the supernatant as an internal control and then the mixture was spun in an SW32Ti over 3 mL 20% sucrose cushion at 24,000 rpm for 2 hours at 4 °C. The tube was inverted and the pellet was dissolved in 100 µL PBS at 4 C for 15 minutes. For Western blot analysis, 12 µl of VLP or 25 µg of cell lysate were added to 4XNuPAGE dye (with B-mercaptoethanol), boiled at 95 °C for 5 min, ran on 4-20% Mini-PROTEAN® TGX™ Precast Protein Gels (BioRad, Cat:4561096) and transferred to a 0.2 µm Nitrocellulose membrane. The membrane was blocked in 10% NFDM and stained with primary antibody: anti-N (Sino Biologicals 40143-MM05, 1:1,000 dilution), anti-S (abcam ab272504, 1:500), anti-GAPDH (Cell Signaling 5174S, 1:1,000), anti-p24 (Sigma SAB3500946, 1:2,000) for O/N at 4 °C. Blots were rinsed with TBS-T three times for 5 minutes each and stained with secondary HRP antibody (Bethyl A90-516P (mouse), A120-201P (rabbit) 1:5,000). Imaged using a chemiluminescence kit (Roche 12015200001, Thermo Scientific™ 34096). Image was captured using ChemiDoc™ Imaging System (Biorad 12003153). Densitometry was done using ImageJ software.

### Innate immunity reporter assays

HEK 293T cells were seeded (60,000 cells) in 48-well plates. 24 h later, cells were transfected with IFN-β promoter-FLuc and SV40-RLuc (25 ng each) reporter plasmids and 5 ng of an inducer plasmid (FLAG-RIG-I, FLAG-MAVS, or FLAG-TBK1) and the M-SARS plasmid (100, 250, or 500 ng) using polyethyleneimine (PEI) (Thermo Fisher Scientific). At 24 h post-transfection, the cells were lysed with passive lysis buffer (Promega Corporation). Luciferase activity was determined using the Dual-Luciferase assay kit (Promega Corporation) following the manufacturer’s instructions, using a Sirius Lumat 9507 luminometer (Berthold Detection Systems GmbH). The protein concentration in cellular lysates was determined by Bradford assay (Bio-Rad Laboratories). Proteins (40 μg) were resolved on a 15% tricine-SDS-polyacrylamide gel and transferred to a polyvinylidene difluoride (PVDF) membrane (Thermo Fisher Scientific). Membranes were blocked (5% skim milk in TBS-Tween 0.1%) and incubated with an anti-M SARS rabbit monoclonal antibody (MA5-46924; Thermo Fisher) at a 1:2500 dilution, an anti-HA mouse monoclonal antibody (H9658; Sigma-Aldrich) at a 1:5000 dilution, an anti-FLAG mouse monoclonal antibody (F1804; Sigma-Aldrich) at a 1:1000 dilution or an anti-GAPDH mouse monoclonal antibody (MA5-15738; Thermo Fisher) at a 1:5000 dilution in TBS-T. As secondary antibodies, we used a goat anti-mouse horseradish peroxidase (HRP)-conjugated antibody or a goat anti-rabbit HRP-conjugated antibody (Merck-Millipore) at a 1:10000 dilution (5% skim milk in TBS-T). Proteins were detected using the SuperSignal West Femto chemiluminescence kits (Thermo Fisher Scientific).

### Anti-Sars-CoV-2 M ELISA

An in-house ELISA was performed based on previously mentioned protocols [28, 52]. Four biotinylated 20-mer peptides spamming the first 20 amino acids of M protein and including the mutations in site 3, and one 18-mer peptide of the N protein amino acids 153-170 (Elim Biopharmaceuticals), were reconstituted in DMSO and working stocks were prepared in water to a final concentration of 10 ug/mL. One ug of each peptide, or phosphate-buffered saline (PBS) as background control, were incubated in streptavidin-coated ELISA plates (Pierce) overnight at 4°C. Plates were 4 times washed with PBS 0,05% Tween-20 and blocked with a commercial blocking buffer (Invitrogen). Thirty-two sera samples from first-wave SARS-CoV-2 infected individuals with high anti-SARS-CoV-2 IgG titers previously determined [3, 30], and a non-SARS-CoV-2 immune sera used as control, were diluted at 1:100 in blocking buffer and added to the plate overnight at 4°C. Plates were washed five times and a horseradish peroxidase-conjugated goat anti-human IgG (KPL) diluted 1:1000 in blocking buffer was added. After 45 min of incubation at room temperature, plates were washed five times and incubated with 100 uL of a tetramethylbenzidine (TMB) substrate solution (Euroimmun) until development of blue color. The reaction was stopped with 100 uL of a Stop solution (Euroimmun) and the optical density at wavelength of 450nm was measured. Values obtained from background controls were subtracted from peptide measurements for each sample, and a cut-off of positivity was determined as two standard deviations from values obtained with the control sera.

### AlphaFold analysis

To assess whether the M-protein homodimers of the WT, BA.1, BA.2, BA.5, and JN.1 variants differ structurally beyond the model-to-model variability intrinsic to AlphaFold 3, we predicted the M homodimer of each variant with the AlphaFold 3 web server (alphafoldserver.com; two copies of the M sequence per job). The server returns five ranked predictions per job, each from an independent random seed; all five were retained, yielding 25 structures (5 variants × 5 models). The aligned models were exported as individual PDB files in PyMOL (open-source v3.1.0). All subsequent analyses used Cα atoms only and were performed in Python 3.13 with Biopython 1.87 (coordinate parsing), NumPy 2.4, and SciPy 1.17.

To estimate the within-variant noise floor, the five models of each variant were superposed by the Kabsch algorithm and the per-residue Cα RMSF and pairwise RMSD were computed. Positions with a mean within-variant RMSF below 3 Å were defined as the structured core (393 of 444 Cα positions; 222 residues per protomer × 2 chains); the 51 excluded positions correspond to the flexible N- and C-termini (RMSF up to ∼20 Å). For each variant a representative model was selected as the medoid (minimum summed core Cα RMSD to the other four), avoiding bias from an arbitrary choice.

For the cross-variant comparison, the five representatives were superposed on the structured core and a 5×5 pairwise Cα RMSD matrix was computed and hierarchically clustered (average linkage). To localize differences per residue, all 25 models were brought into a common frame by iterative superposition on the core, and an ANOVA-style F-ratio (between-variant variance / within-variant variance; df = 4 and 20) was computed for each Cα position, with p-values corrected across positions by the Benjamini–Hochberg procedure (significance at q < 0.05). Finally, the per-residue between-variant Cα deviation was written to the B-factor column of the WT representative and rendered on the structure.

## Acknowledgements

This work was supported by the Agencia Nacional de Investigación y Desarrollo ANID Chile (FONDECYT 11261435, 3220310, and SENTINET CIN250062 to Barrera-Vásquez A., FONDECYT 1230718 to Ramos H. and Angulo J., and doctoral fellowship 21230810 to Ramos H.). M.O. received support from the Roddenberry Foundation, James B. Pendleton Charitable Trust, and P. and E. Taft, and is supported by the Gladstone Institutes. M.O. is a Biohub San Francisco Investigator and the Nick and Sue Hellmann Distinguished Professor. We thank Dr. Nicole Le Core and Dr. Elvira Balcells for sharing anonymized human sera samples from projects ANID COVID0920, Fondo de Adopción Tecnológica SiEmpre, SOFOFA Hub, and Ministerio de Ciencia, Tecnología, Conocimiento e Innovación, Chile. We thank Dr. Françoise Chanut for feedback on the manuscript.

