## Supplementary Information for "Epistasis between SARS-CoV-2 M and N Proteins Balances Particle Assembly and Immune Evasion"

**Short title:** M-N Epistasis Balances Assembly-Immunity Tradeoffs

**Authors**: Aldo Barrera-Vasquez[1,2], Mir M. Khalid[3], Hade Ramos[1,4], Julia Rosecrans[3], Marcela Ferres[1,2], Jenniffer Angulo[1,4], Melanie Ott[3,5,6]*, Taha Y. Taha[3,7]*

**Affiliations**:
[1] Departamento de Enfermedades Infecciosas e Inmunología Pediátricas. Escuela de Medicina. Pontificia Universidad Católica de Chile, Santiago, Chile.
[2] SENTINET, Santiago, Chile
[3] Gladstone Institutes, San Francisco, CA, USA
[4] Facultad de Ciencias Biológicas. Pontificia Universidad Católica de Chile, Santiago, Chile.
[5] Department of Medicine, University of California, San Francisco, CA, USA
[6] Chan Zuckerberg Biohub – San Francisco, San Francisco, CA, USA
[7] Department of Bioengineering and Therapeutic Sciences, University of California, San Francisco, CA, USA

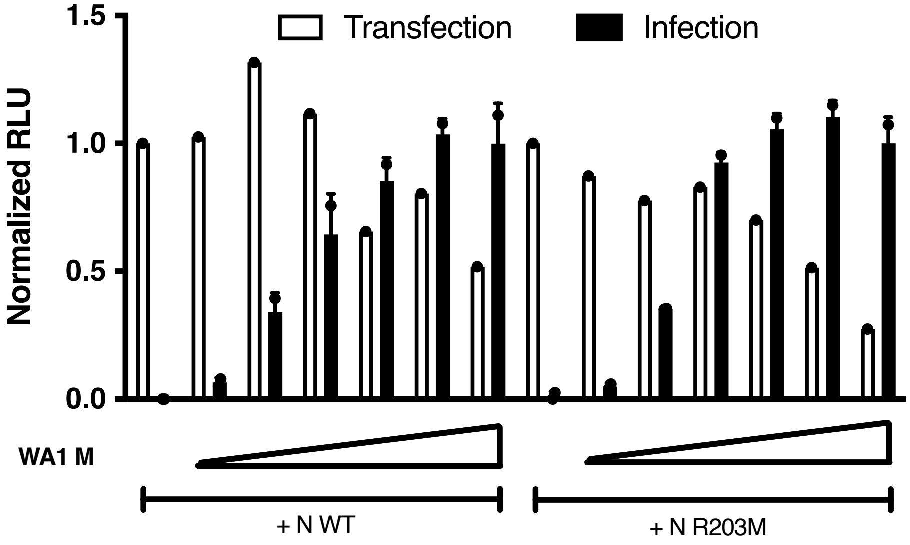


**Supplementary Figure 1. Robust particle production depletes reporter expression in transfected cells.** Luciferase readout of transfected BHK-21 cells and infected VAT cells with varying amounts of M expression vector and either WT or R203M N protein. Data are presented as mean +/- SD of two technical replicates from a single transfection experiment and normalized to the 0 μg M protein control.

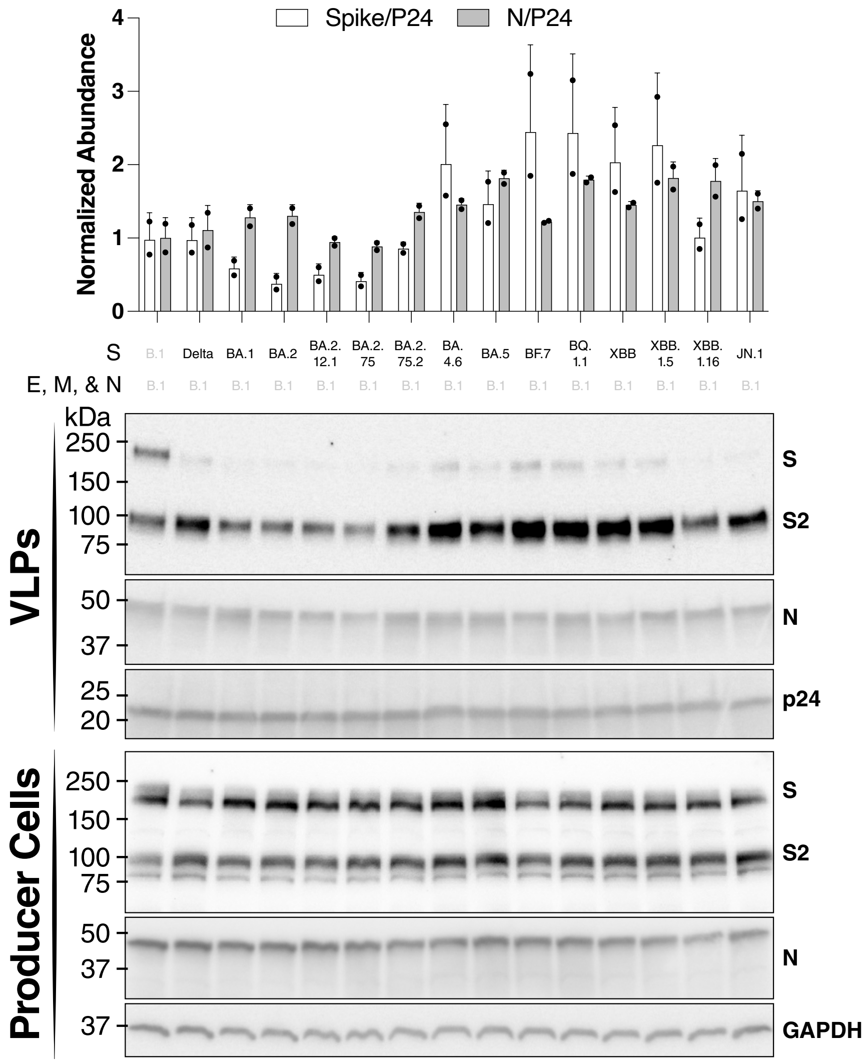


**Supplementary Figure 2. Spike protein variants do not impact VLP production.** Western blot analysis of VLP producer cells and purified VLPs for the Spike and N proteins. GAPDH was used as a cellular internal control, and p24 was used as VLP purification internal control for spike-in lentiviral particles. The composition of the VLPs is shown on top of the blots. Blots are representative of two independent biological replicates. The top panel shows measured VLP abundance normalized by p24 abundance and presented as mean +/- SD deviation. *, $p < 0.05$ using two-sided Student's T-test.

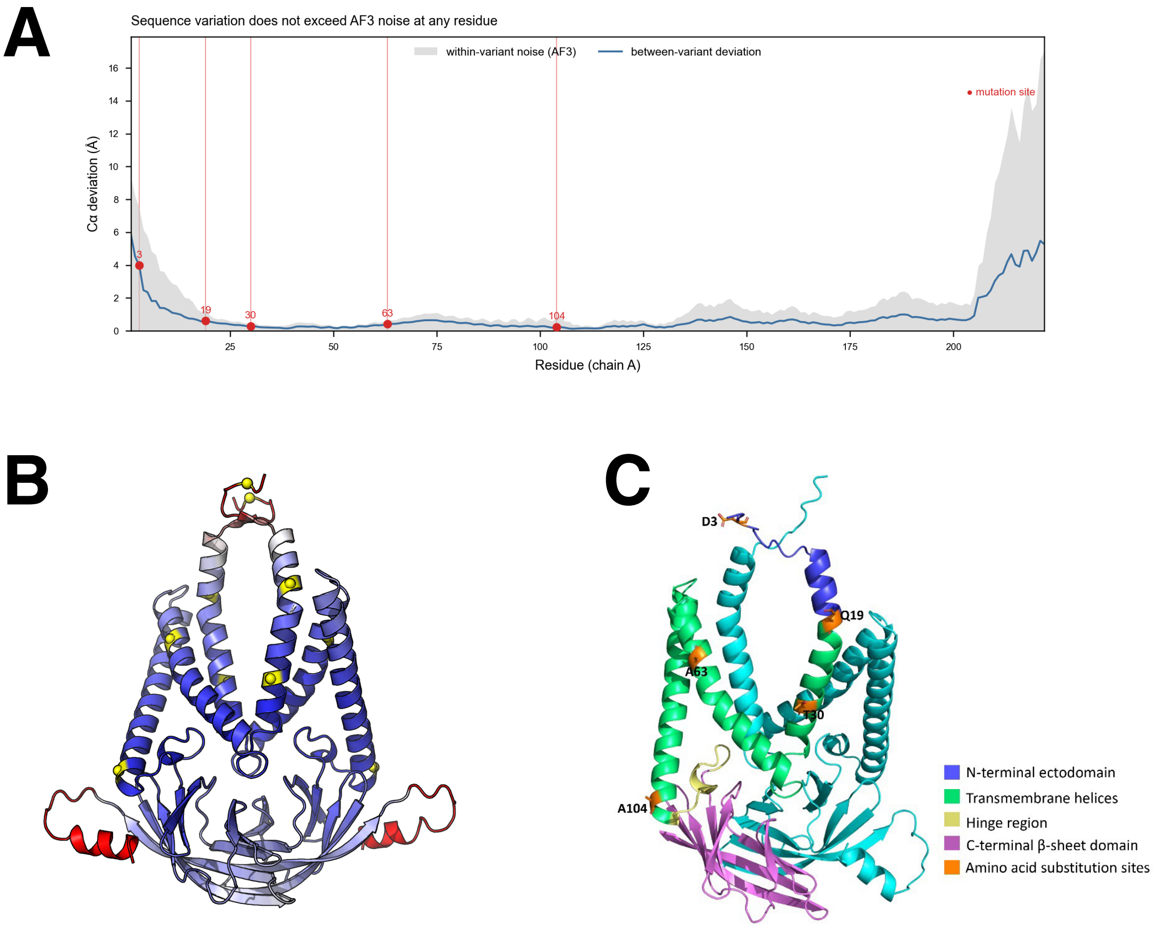


**Supplementary Figure 3. M mutations do not significantly alter the protein's structure.** A) The M protein structure of WT, BA.1, BA.2, BA.5, and JN.1 were predicted using AlphaFold 3 and the between model variance plotted per residue (see methods). The mutations along the protein sequence are indicated with red dots. **B)** The variance between models in A is rendered on M protein structure using Pymol. Blue color shows low variance and red color shows high variance. Yellow colored residues are those where mutations occurred (see methods). **C)** The M protein structure was predicted using AlphaFold 3, showing a moderate confidence range (pLDDT ~50–70), and aligned to the available experimental structure (PDB: 8CTK), yielding an RMSD of ~3 Å (the predicted full-length model was used due to missing residues in the crystallographic structure). The figure depicts the M protein as a dimer, with one monomer shown in cyan and the second monomer colored by structural domains: the N-terminal ectodomain in blue, the transmembrane helices in green, the hinge region in yellow, and the C-terminal β-sheet domain in pink. Residues corresponding to the positions of the amino acid substitutions analyzed in this study are shown as orange sticks and labeled with their residue identity and position. The structure was visualized using PyMOL.

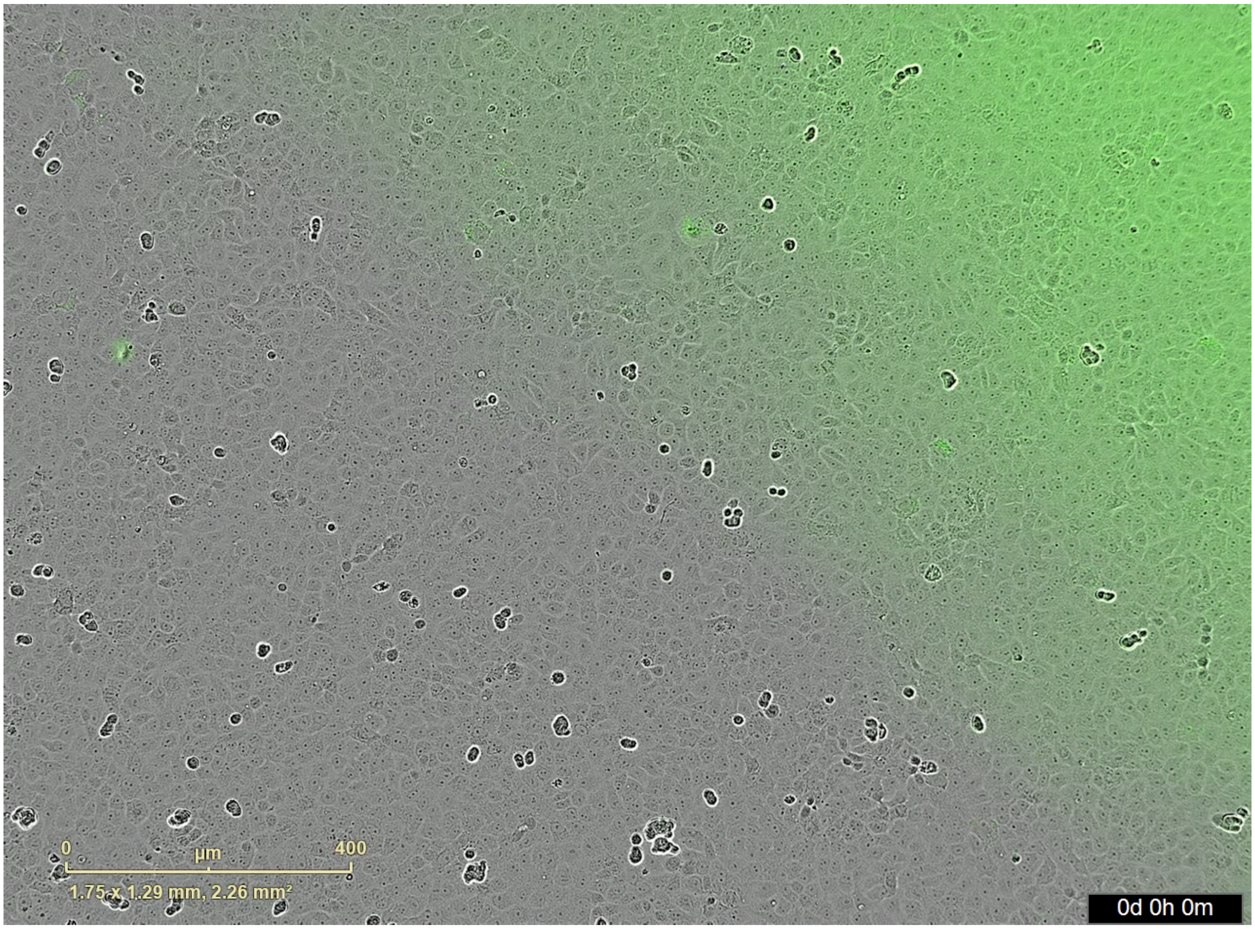


**Supplementary Video S1. SARS-CoV-2 M replicon shows cell-cell spread during infection.** Live cell imaging and fluorescence readout of infected VAT cells over 5 days post-infection. The video is representative of three independent biological replicates. The video was produced on an Incucyte system.
